# Engineered gut symbionts mediate cross-phylum antagonism to suppress uropathogenic *Escherichia coli* colonization

**DOI:** 10.64898/2026.05.11.724322

**Authors:** Jay Fuerte-Stone, Joyce Ghali, Sandra Valaitis, Mark Mimee

**Affiliations:** Committee on Microbiology, University of Chicago, Chicago, IL 60637, USA; Committee on Molecular Metabolism and Nutrition, University of Chicago, Chicago, IL 60637, USA; Urogynocology, University of Chicago Medicine, Chicago, IL 60637, USA; Department of Obstetrics and Gynecology, Section of Urogynecology, University of Chicago, Chicago, IL 60637, USA; Department of Microbiology, University of Chicago, Chicago, IL 60637, USA; Pritzker School of Molecular Engineering, University of Chicago, Chicago, IL 60637, USA; UChicago Medicine AdventHealth Medical Group Urogynecology, Hinsdale, IL 60521, USA

**Author notes:** **Corresponding author:** Mark Mimee, **Email:**. **Author Contributions:** J.F.S. and J.G. designed and performed experiments. S.V. provided clinical isolates. J.F.S. and M.M. conceived this study, analyzed data, discussed results, and wrote the manuscript. **Competing Interest Statement:** The authors declare no competing interests.

**Keywords:** Microbiome, synthetic biology, bacterial antagonism, uropathogenic *Escherichia coli*

## Abstract

Urinary tract infections (UTIs) are among the most common bacterial infections globally and create a large burden on the healthcare system. Uropathogenic *Escherichia coli* (UPEC) account for the majority of UTIs and increase the risk of recurrence. The standard treatment is antibiotics and, with the rise of multi-drug resistant UPEC lineages, there is a need for alternative treatments and prevention. Colicins, bacteriocins targeting and produced by *E. coli*, have previously been shown to inhibit the growth of pathogenic *E. coli* and are a promising alternative. Here, we engineer commensal Bacteroidaceae to secrete colicins via outer membrane vesicle (OMV) targeting signal peptides to suppress *E. coli* in the mouse gut. Secreted colicins were assessed for their ability to kill primary clinical isolate UPEC strains, including epidemic multi-drug resistant ST131 strains, along with other pathogenic and type strains. Specifically, secreted colicin E7, from *Phocaeicola vulgatus* fully eliminated of several UPEC strains in culture. In mice, *P. vulgatus* secreting colicin E7 prevented the extended colonization of two clinical UPEC strains and restored microbiome diversity. Together, this work shows the viability of secreted, heterologous antimicrobials from *P. vulgatus* as prophylactic treatment against the colonization of pathogenic *E. coli* utilizing cross-phylum antagonism in the gut.

**Significance Statement:** Recurrent urinary tract infections can be driven by intestinal reservoirs of uropathogenic *Escherichia coli* that are difficult to eliminate and increasingly recalcitrant to conventional antibiotic therapy. Here, we show that engineered gut symbionts from the Bacteroidaceae family can secrete targeted protein antibiotics to selectively kill these uropathogenic *E. coli*. Leveraging outer membrane vesicle-based secretion, we demonstrate that bacteriocin secretion can prevent gut colonization by clinically relevant pathogens, while preserving overall microbiome diversity. This work establishes a strategy for programmable, cross-phylum antimicrobial delivery within the gut microbiome, providing a potential alternative to conventional antibiotics for preventing recurrent infections and other enteric diseases.

## Introduction

Urinary tract infections (UTIs) are among the most common bacterial infections globally, primarily affecting people with vulva and placing a large burden on healthcare systems [1,2]. The most common cause of UTIs is uropathogenic *E. coli* (UPEC), which account for approximately 80% of uncomplicated cases [1,3]. UTIs have a high likelihood of recurrence, with 27% of female patients reporting a second UTI within six months of the first infection [4] and an increased risk of recurrence for *E. coli* infections [5]. Reinfection by the same strain is seen in 50% of recurrent UTIs, suggesting persistence of pathogens in either the urogenital tract or the intestine after treatment [6]. The gut can act as a primary reservoir for UTI pathogens, with UTIs occurring from transfer of gut-resident pathogens to the urethra and subsequent colonization of the bladder [1]. UPEC strains have been shown to asymptomatically colonize both the intestine and urinary tract and antibiotic-induced blooms of gut *E. coli* in the gut and decreased microbiome diversity correlating with urinary tract colonization[7]. Antibiotic treatment is standard, with prophylactic antibiotic treatment often prescribed for recurring UTIs [8]. There is a growing concern surrounding extended-spectrum beta-lactamases (ESBL) producing *E. coli*, as well as fluoroquinolone resistance, observed in patient UPEC isolates [9]. The most common and globally distributed UPEC clone, pandemic lineage ST131, is increasingly associated with these types of antibiotic resistance [10]. There is therefore a need to develop alternative treatment options that do not rely on conventional antibiotics and are able to prevent persistence of UPEC in the gut.

Antagonistic systems are abundant in the gut microbiome, allowing for bacterial strains to compete within the community [11]. Many of these systems target close taxonomic relatives to the producer strain, as close relatives tend to occupy similar physical and nutritional niches [11]. Bacteriocins are diffusible, protein-based antimicrobials that typically exhibit a narrow target range and have been proposed as an alternative to antibiotics due to both targeted killing and high potency [12,13]. *E. coli*-produced and targeting bacteriocins, or colicins, are encoded on plasmids with an immunity protein and are tightly regulated by repressor LexA, which is inactivated by the SOS response [14]. Colicins are made up of three domains: N-terminal domain for membrane translocation, receptor binding at the center domain, and an active site at the C-terminal domain [15]. The membrane receptor and mechanism of action vary for each colicin. Cell entry is mediated by binding to a nutrient-related membrane protein, such as class E colicins binding to vitamin B_12_ receptor BtuB [16]. Membrane translocation is then dependent on either the Ton or Tol systems [16]. Colicins kill target cells through pore formation, nuclease activity, or peptidoglycan biosynthesis inhibition [15].

Colicins and colicin-like proteins, such as pyocins and klebicins, have been proposed as alternatives to antibiotics for the control of pathogens. Previous studies have shown inhibition of *Pseudomonas aeruginosa* by *E. coli* engineered to produce and release pyocins via secretion [17] and lysis [18]. Additionally, lytic bacteriophage engineered with colicins or klebicins exhibited increased killing and reduce resistance both in vitro and ex vivo in UTI patient urine samples [19]. Here, we propose utilizing colicin as a method for preventing gut colonization of opportunistic pathogens, such as UPEC. We engineered commensal Bacteroidaceae to constitutively produce and secrete colicins, leveraging the gut as a bioreactor for continuous production in the known UPEC reservoir. We show that secreted colicin is active against a variety of pathogenic *E. coli*, including patient isolated UPEC, and is capable of preventing colonization of UPEC strains in the mouse gut, demonstrating engineered cross-phylum antagonism for pathogen resistance.

## Results

### Genomic analysis of uropathogenic *E. coli* patient isolates

We obtained 25 *E. coli* strains isolated from patients at the University of Chicago who presented with recurrent urinary tract infections (UTI), as defined by two or more infections in 6 months. We analyzed the genomes of these UPEC strains, along with 9 other pathogenic and nonpathogenic *E. coli* and commensal *E. coli* Nissle 1917 (Figure 1). We identified 14 multilocus sequencing types (MLSTs) from the 25 clinical samples. The most abundant were ST131 (4 strains), ST 69 (4 strains), and ST95 (3 strains) (Table 1). These three MLSTs, along with ST73, are the most prevalent extraintestinal pathogenic *E. coli* (ExPEC) sequence types, including those isolated from UTIs [20,21]. Additionally, these isolates also represent 3 phylotypes: B_1_ (2 isolates), A (3 isolates), and B_2_ (16 isolates). Group B_2_ is the most prevalent phylotype in clinical UPEC isolates [22–24] and have a high capacity for biofilm formation [1,25]. Biofilm formation is a key mechanism in recurrent UTIs, allowing for host defense evasion and preventing antibiotic diffusion [26].

**Figure 1.**
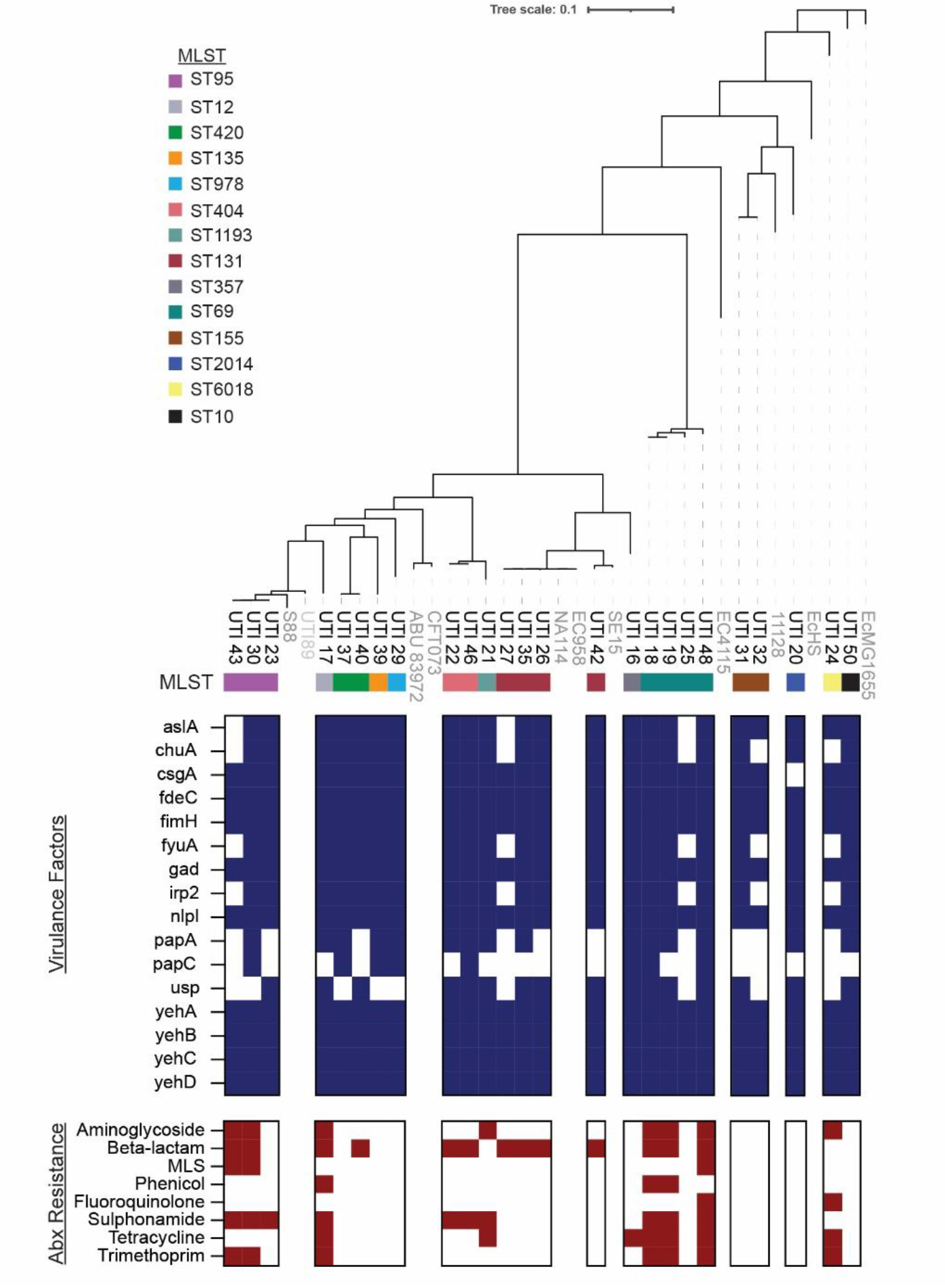
Primary clinical isolate UPEC strains harbor virulence factors for colonization of the urogenital tract and antibiotic resistance markers. 25 *E. coli* strains were isolated from patients with recurrent urinary tract infections (rUTIs). Phylogenetic distribution (top) of UTI strains (black text) and 10 other pathogenic and laboratory E. coli strains (gray text) were determined by constructing a pan-genome and core genome alignment. Multilocus sequence type (MLST) was determined for each UPEC isolate from this study (MLST 2.0, Achtman scheme). Virulence factors (middle) were determined for each strain, with blue squares indicating presence of a given gene. Antibiotic resistance was also identified computationally, with maroon squares indicating resistance for a given antibiotic. (MLS = Macrolides, lincosamides, streptogramins).

Virulence factor analysis showed that all UPEC strains harbored *fdeC, fimH, nlpI*, and *yehABCD*, all genes associated with adherence to epithelial cells and formation of biofilms [27–29]. Additionally, all UPEC strains harbor a gene for glutamate decarboxylase *gad* which confers resistance in acidic environments [30], and tellurite resistance gene *terC* [31]. The majority of isolates also have genes for adhesion (*papA, csgA*) [32,33], tissue invasion *aslA* [34], and iron uptake and utilization (*chuA, fyuA, irp2*) [27]. During infection, iron is sequestered by the host [35] and UPEC strains have been shown to both harbor a variety of iron acquisition pathways and up-regulate iron acquisition genes during infection [36,37]. 16 UPEC isolates harbor *usp* which encodes a genotoxin active against epithelial cells which has been shown to be present in a majority of UPEC strains, compared to around a quarter of feces isolates [38,39]. Together, this genetic analysis shows the virulence of these UPEC isolates and confirms findings from previous studies on the widespread biofilm formation and iron utilization genes in uropathogenic isolates. Additionally, the diversity in phylogeny, sequence type, and virulence determinants supports this strain panel as a clinically relevant system for testing novel antimicrobials targeting UPEC.

### Engineering colicin-secreting Bacteroidaceae

As UTIs often begin with the transit of *E. coli* from the gut to the urethra [1], we aimed to design a method to prevent colonization of the gut with uropathogenic strains. Bacteroidaceae are dominant members of the human gut microbiome [40] and are largely commensal, playing a key role in the microbiome as primary polysaccharide consumers, production of short chain fatty acids (SCFAs), immunoregulation, and for their role in pathogen suppression in the gut [41,42]. Their role as potential live biotherapeutics and advances in synthetic biology that enable programmable secretion of heterologous proteins [43] makes Bacteroidaceae members an ideal chassis candidate for a antimicrobial secretion system. Lipoproteins are naturally enriched on *Bacteroides* outer membrane vesicles (OMVs) and their signal peptides have been shown to target fused heterologous cargo to OMVs, resulting in both OMV-attached and freely soluble proteins in supernatant [43,44]. Due to the necessity of both the N- and C-termini for colicin translocation and mechanism of action [15], freely soluble colicin is needed for suppression of *E. coli* (Figure 2A). Using these signal peptides, we designed 12 constructs for the secretion of colicins from Bacteroidaceae (Figure 2B). For each construct, an immunity protein was placed under constitutive promoter P_BfP1E6_ and followed by the corresponding colicin fused to a signal peptide with a (Gly_4_Ser)_2_ flexible linker. Three colicins (Colicin M, E6, and E7) were chosen for varied modes of action and efficacy in antagonism of pathogenic *E. coli* [45]. Colicin M (ColM) inhibits peptidolglycan biosynthesis by hydrolyzing lipid II, leading to cell lysis [46,47], colicin E6 is a 16S rRNase, inhibiting translation [48], and colicin E7 exhibits nonspecific DNase activity [49]. Constructs were then integrated into the genomes of *Bacteroides fragilis, Bacteroides thetaiotaomicron, Bacteroides uniformis*, and *Phocaeicola vulgatus* using the pNBU1 integration system. Integrated strains will be referred to as ‘species (signal peptide)-colicin’, for example Pv BT_1491-CM refers to *P. vulgatus* with colicin M fused to signal peptide BT_1491.

**Figure 2.**
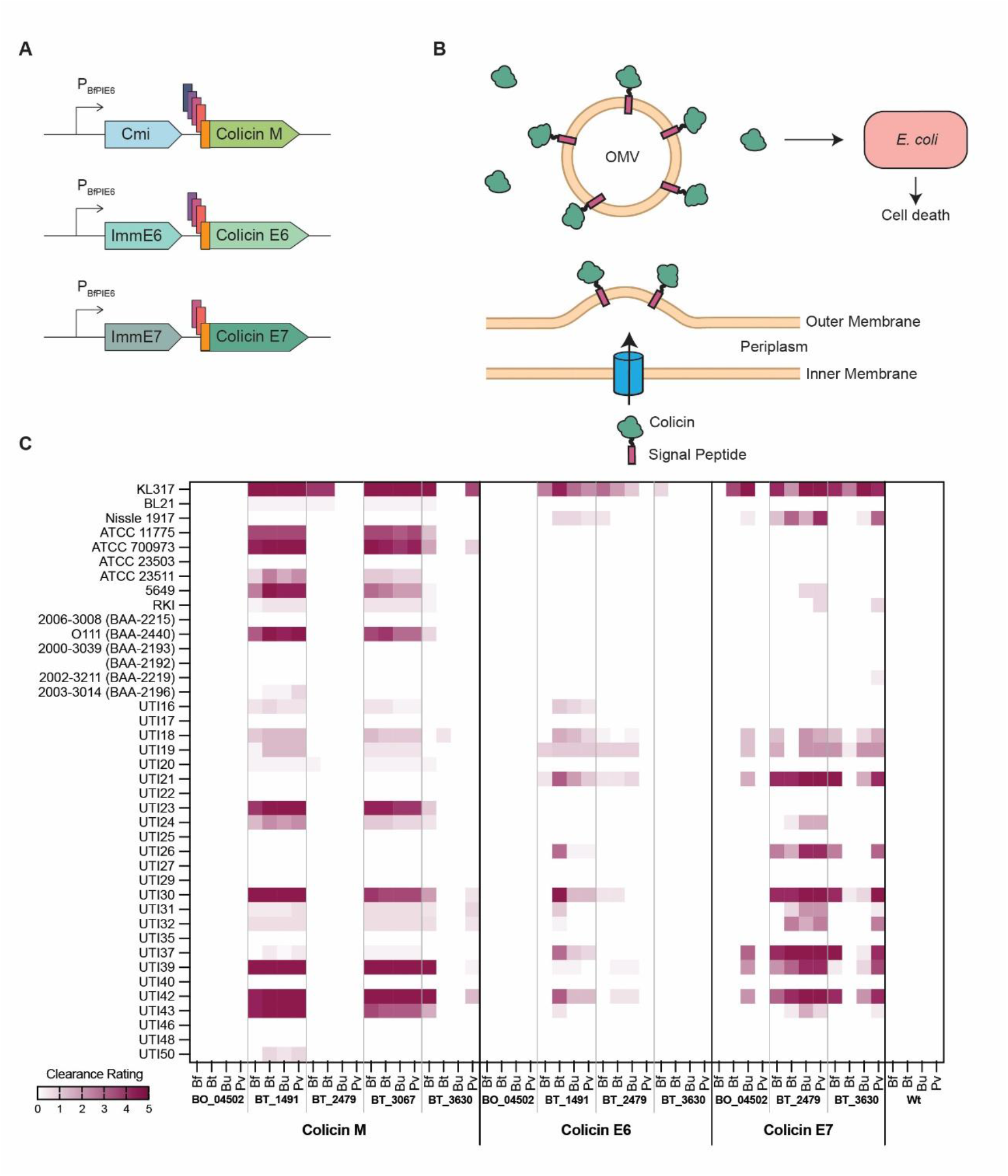
UPEC isolates are susceptible to colicins secreted by engineered Bacteroidaceae. (A) Operon design for colicin secretion system. Colicins were fused to 3-5 signal peptides and placed under constitutive promoter P_BfP1E6_ following the appropriate immunity protein. Immunity protein-colicin pairs were (top) colicin M immunity protein (Cmi) and colicin M, (middle) E6 immunity protein (ImmE6) and colicin E6, and (bottom) E7 immunity protein (ImmE7) and colicin E7. Operons were placed in plasmid pNBU1 for integration into *B. fragilis* (Bf), *B. thetaiotaomicron* (Bt), *B. uniformis* (Bu), and *P. vulgatus* (Pv). (B) Illustrative diagram of secretion of colicins via outer membrane vesicles (OMVs) from engineered Bacteroidaceae strains resulting in colicins attached to OMVs and free in solution, resulting in cell death of *E. coli*. (C) Heatmap of the susceptibility of 40 *E. coli* strains to secreted colicins from colicin-producing Bacteroidaceae strains, as determined by soft agar overlay assays. Clearance rating was based on appearance for each spot as follows, with darker shading corresponding to increased clearance: no visible clearance (0), lightened area (1), hazy area (2), clearance smaller than original Bacteroidaceae spot (3), clearance equal to original spot (4), and clearance larger than original spot (5). Values represent mean of three replicates.

### In vitro testing of Bacteroidaceae-secreted colicins against UPEC isolates

To evaluate the killing ability of these secreted colicins, we performed soft agar overlay assays. Cultures of Bacteroidaceae strains harboring the colicin constructs and wild-type controls were spotted on agar plates and allowed to grow overnight. Spots were then physically removed and remaining cells killed with chloroform vapors. The remaining colicin represents what had been secreted into the agar during bacterial growth. Plates were then covered in soft agar containing *E. coli* of interest. We tested 48 colicin-producing Bacteroidaceae strains, along with four wild-type controls, against 40 *E. coli* strains and scored clearance on a scale of 0-5 with 5 being full *E. coli* clearance (Supplementary Figure 2A). *E. coli* strains in this screen included nonpathogenic type strains, the UPEC patient isolates, pathogenic isolates from a variety of sources, and strains representing the “big six” designation of non-O157 serogroups causing foodborne illness [50]. While individual peptide-colicin pairs tended to behave similarly across species, the success of secretion peptide-colicin pairs was more idiosyncratic (Figure 2C). We observed that strains utilizing signal peptide BO04502 exhibited no clearing, possibly due to lack efficient colicin secretion. Colicin E7 demonstrated effective clearance when fused with BT_2479 and BT_3630, whereas colicin M favored BT_1491 and BT_3067. Additionally, colicin E6 showed primarily weak clearance against most *E. coli* stains and therefore was not included in the rest of this study. The colicin N-terminus is responsible for initiating entry into the target cell [15] and it is therefore possible that fusion of the signal peptide to the N-terminus disrupted this interaction and prevented efficient entry into the cell. BT_1491-CM and BT_3067-CM exhibited high clearance of multiple *E. coli* strains, including 5 UPEC isolates and O111 (BAA-2440), a Shiga toxin-producing E. coli (STEC) which represents one of the “big six” serogroups causing food-borne illness [50].Additionally, all three ST95 strains were seen to be highly susceptible to secreted ColM. BT_2479-E7 and BT-3630-E7 showed high clearance of 6 UTI strains but no other pathogenic strains. Interestingly, secretion of BT_3630-E7 was extremely weak, as clearance was only moderately seen in KL317. Based on these results, we chose to move forward with BT_1491-Bt and BT_2479-E7 for further in vitro testing.

Based on the agar overlay assay results (Supplementary Figure 2B-F), we next selected KL317, a laboratory *E. coli* K-12 strain, and four UPEC isolates (UTI21, UTI30, UTI37, and UTI42) to evaluate the killing capacity of BT_1491-CM and BT_2479-E7 supernatants. OD_600_-normalized and filtered supernatant from cultures of Bt BT_1491-CM, Pv BT_2479-E7, wild-type, or PBS was mixed 1:1 with cultures of *E. coli* at ∼10^6^ CFU/mL. Cultures were sampled at 0, 2, 4 and 24 hours to determine the titer of viable cells. Three *E. coli* strains (KL317, UTI30, and UTI42) were susceptible to both ColM and E7 in the plate assay and for these strains we tested supernatants both individually and in combination. For all groups, KL317 had an initial reduction in titer, followed by a resurgence in population, though the mixed colicin supernatant suppressed this growth more than the individual supernatants (Figure 3). No *E. coli* strain was fully suppressed by Bt BT_1491-CM supernatant, with UTI30 titers only dropping tenfold (Figure 3 A, C, and E). UTI21 and UTI30 were initially killed to undetectable levels in the first two hours by Pv BT_2479-E7 supernatant, but growth rebounded by 24 hours (Figure 3B and C). Pv BT_2479-E7 supernatant reduced UTI37 and UTI42 to undetectable levels by 2 hours, with no growth seen at 24 hours (Figure 3D and E). Additionally, UTI30 and UTI42 cultures were fully killed by the combination supernatant by 2 hours and remained undetectable (Figure 3C and E). Together, we show that BT_2479-E7 is able to kill UPEC strains in culture and prevent resurgence both alone and in combination with BT_1491-CM.

**Figure 3.**
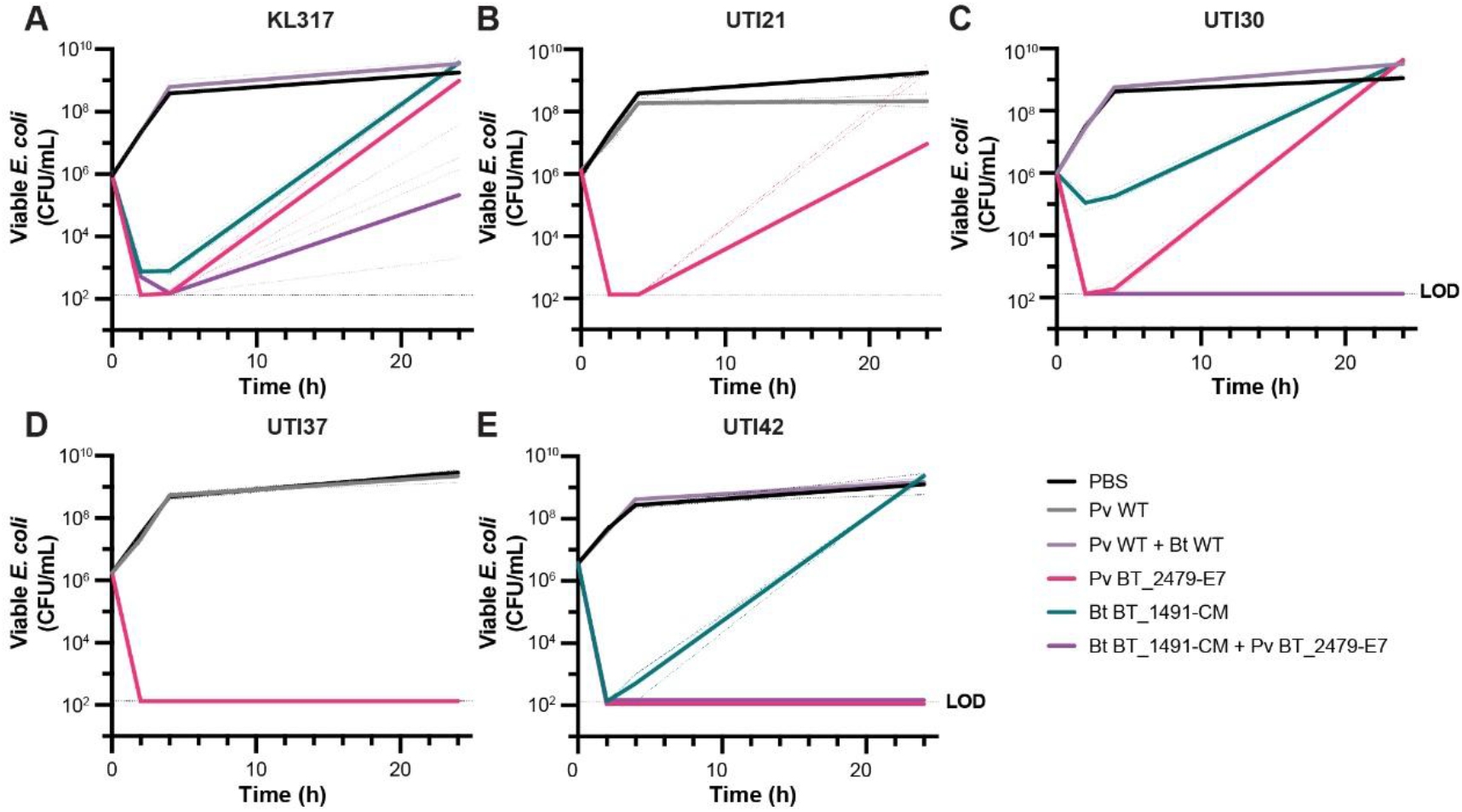
Colicin E7 secreted by P. vulgatus and colicin M secreted by B. thetaiotaomicron kill UPEC isolates in culture. 10^6^ CFU/mL of *E. coli* was incubated with an equal volume of filtered supernatant from either Pv BT_2479-E7 (pink), Bt BT_1491-CM (teal), or both (purple). Viable *E. coli* was enumerated by CFU plating at 0, 2, 4, and 24 hours for (A) KL317, (B) UTI21, (C) UTI30, (D) UTI37, and (E) UTI 42. Solid lines represent geometric means of three replicates (individual replicates in dashed lines).

### Pv BT_2479-E7 prevents colonization of UPEC and restores diversity in a mouse gut

We next evaluated Pv BT_2479-E7 as a method of preventing colonization of UPEC in the mouse gut. A prophylactic infection model was chosen to emulate prevention of recurrent UTIs post-antibiotic treatment of an initial infection. Additionally, we used C57BL/6 mice from Jackson Laboratory because these mice lack native Enterobacteriaceae [51], allowing for facile gut colonization of human *E. coli* strains and CFU enumeration on MacConkey plates, without contamination of native Gram negative facultative anaerobes. We confirmed no bacterial growth on MacConkey plates when untreated mouse fecal samples were incubated aerobically for 48 hours at 37C. Mice were colonized with either BT_2479-E7 or wild-type *P. vulgatus* and, after streptomycin treatment and a washout period, gavaged with a single dose of ∼5x10^4^ CFU of either UTI37 or UTI42 (Figure 4A). Fecal samples were collected over the course of 7 days and processed for both viable *E. coli* and total microbial DNA. For both UPEC strains, colonization was rapid in mice with wild-type *P. vulgatus*, with at least 10^8^ CFU/g feces reached for all but one mouse (Figure 4B and F). UTI42 was not detected in any Pv BT_2479-E7 colonized mice over the course of the experiment, indicating complete prevention of colonization of the UPEC strain. UTI37 was detected at low levels in two mice colonized with Pv BT_2479-E7 on Day 2, one on Day 4, and was undetectable in all mice by Day 7. Overall, colicin-secreting *P. vulgatus* can effectively suppress colonization of UPEC strains in the mouse gut.

**Figure 4.**
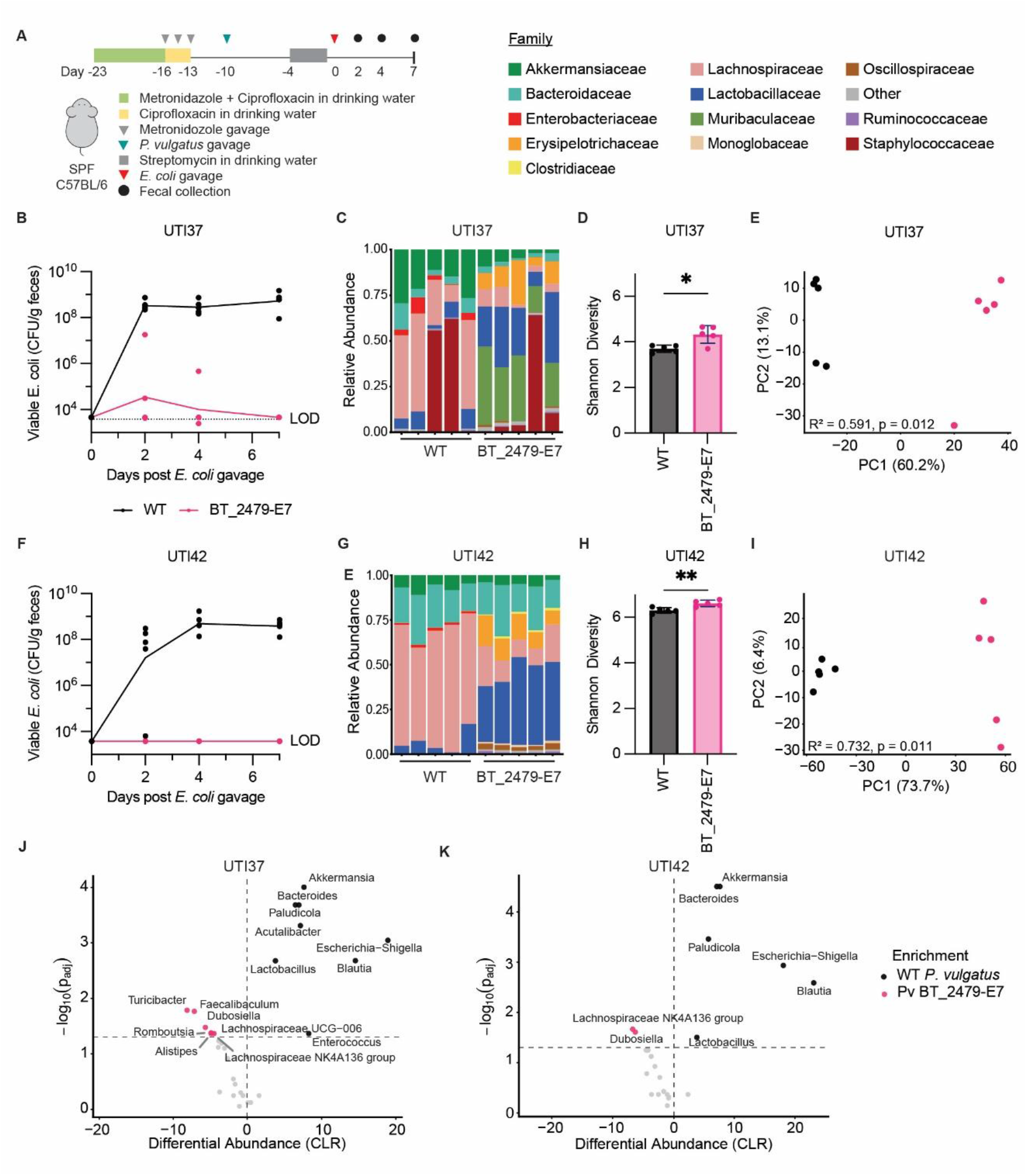
Pv BT_1491-E7 prevents the colonization of primary patient UPEC isolates in mice and restores diversity of the microbiome. (A) SPF C57BL/6 were treated with metronidazole and ciprofloxacin for ten days and then colonized with either wildtype *P. vulgatus* or Pv BT_2479-E7. Mice were then treated with streptomycin for three days and gavaged with either UTI37 (B-E, J) or UTI42 (F-I, K) (n=5). (B and F) *E. coli* burden in mouse fecal pellets, as enumerated by CFU on selective plates. (C and G) 16S rRNA profiling of bacterial communities at Day 7 at the family level. (D and H) Alpha diversity analysis of Day 7 samples comparing groups colonized with wildtype *P. vulgatus* or Pv BT_2479-E7, quantified using Shannon diversity index. Statistical significance determined by unpaired t-test with Welch’s correction (*p < 0.05, **p < 0.01). (E and I) Principle coordinate analysis of Aitchison distances (Eucledian distance on centered log-ratio-transformed counts) showing separation of microbial community composition by wildtype *P. vulgatus* or Pv BT_2479-E7 colonization. Differences in community composition were assessed by permutational multivariate analysis of variance (PERMANOVA), with the R^2^ and P value shown on the plot. (J and K) Volcano plots showing differential abundance analysis performed using ALDEx2, with each point representing a taxon plotted by centered log-ratio (CLR) effect size (x-axis) and −log_10_ false discovery rate (FDR-adjusted P value; y-axis). Positive effect sizes indicate enrichment in the Pv BT_2479-E7 colonized mice group and negative values indicate enrichment in the wild-type colonized group. Genera with FDR < 0.05 are highlighted.

To evaluate the ecological consequences of Pv BT2479-E7, we profiled microbiome composition using 16S rRNA sequencing (Figure 4C and 4G). In both UTI37 and UTI42 infected groups, colonization with Pv BT_2479-E7 resulted in a significant increase in alpha diversity compared to wildtype colonized groups (p=0.019 and p=0.005, respectively) (Figure 4D and 4H). Principle coordinate analysis showed clear separation by treatment groups for both UTI37 and UTI42 infected mice as well (PERMANOVA: R^2^ = 0.591, p = 0.012; R^2^ = 0.732, p = 0.011) (Figure 4E and I), indicating a substantial shift in overall community composition. In UTI37-infected mice, Pv BT_2479-E7 colonization was associated with enrichment of multiple genera (*Alistipes, Dubosiella, Faecalibaculum, Romboutsia, Turicibacter*, and members of Lachnospiraceae), wildtype *P. vulgatus* colonization was associated with enrichment of *Escherichia, Blautia, Akkermansia, Bacteroides*, and *Enterococcus* (Figure 4J). In constrast, UTI42-infected mice exhibited fewer differentially enriched taxa, with Pv BT_2479-E7 associated with Lachnospiraceae NK4A136 and *Dubosiella*), while wildtype-colonized mice were enriched for *Escherichia*, Blautia, *Akkermansia*, and *Bacteroides* (Figure 4K). Together, these results show that colicin E7 secreted from engineered *P. vulgatus* prevents the colonization of UPEC in the mouse gut and a shift toward a more diverse gut microbial community.

## Discussion

Here, we engineered strains of Bacteroidaceae to secrete OMV-associated colicins to prevent gut colonization of uropathogenic *E. coli*. We assessed three colicins, ColM, E6, and E7, fused to an array of signal peptides which had previously been shown to enable to heterologous protein secretion from *Bacteroides* [43,44]. Secreted ColM and E7 suppressed the growth of pathogenic *E. coli* in vitro, with BT_2479-E7 preventing rebound growth across several UPEC strains. Additionally, we demonstrated that BT_2479-E7 secreted from *P. vulgatus* prevented colonization by clinically relevant UPEC strains. Notably, UTI42 belongs to sequence type 131 (ST131), the most prevalent UPEC lineage associated with both multi-drug resistance (MDR) and community spread [20,52,53]. Pv BT_2479-E7 fully prevented colonization of UTI42, with the strain remaining undetectable throughout the experiment. Similarly, UTI37 colonization was reduced, with initial partial colonization followed by clearance by Day 7. Together, these results demonstrate that secreted colicin E7 by *P. vulgatus* is sufficient to prevent gut colonization by UPEC, establishing a strategy for engineered cross-phyla antagonism in the gut.

This work presents a potential alternative or adjunct treatment to prophylactic antibiotic use for recurrent UTIs. Longitudinal studies have shown that gut-resident UPEC strains can shift over time and that antibiotic treatment may facilitate colonization or expansion of new strains [54,55]. In these patients, engineered probiotics might be used alongside standard antibiotic treatment to prevent new UPEC colonization and control blooms. Additionally, Aziz et al found that 20% of UTI isolates are due to zoonotic UPEC strains [56]. For patients predisposed to recurrent UTIs, especially those on prophylactic antibiotic treatment, prophylactic treatment with engineered commensals targeting *E. coli* could add an additional preventative measure. More broadly, this approach may be applicable to other enteric infections. Enterotoxigenic *E. coli* (ETEC), a leading cause of traveler’s diarrhea [57] could similarly be targeted through engineered live biotherapeutics to prevent infection. Given the diversity of bacteriocins against gut-resident opportunistic pathogens, including *Acinetobacter* [58], *Klebsiella*, and *Enterococcus*, this strategy may be broadly applicable for targeted removal of pathogens from the gut microbiome.

As with all antimicrobial strategies, the development of resistance is inevitable and must be considered during therapeutic development. Colicins and their immunity proteins are often plasmid-borne and may be already encoded by targeted pathogens or acquired through horizontal gene transfer (HGT) [15]. For example, in our initial screen, several of the rUTI isolates contained colicin E7 and its cognate immunity protein and were therefore resistant to colicin E7 secreting Bacteroidaceae (Supplemental Figure 1A). To mitigate this risk, future approaches may adopt a multi-colicin system by either secreting multiple colicins from a single engineered strain or across a consortium of strains. Given that over 20 colicins have been characterized [14], an expanded colicin repertoire may further enhance efficacy and reduce the likelihood of resistance. More broadly, this work establishes a framework for the delivery of heterologous proteins via Bacteroidaceae-secreted OMVs, enabling applications beyond colicins to other therapeutic payloads in the gut.

## Materials and Methods

### Strains and Culture Conditions

All strains used in this study are listed in Supplementary Table 1. Deidentified primary clinical isolates were streaked from urine samples obtained from patients being treated for recurrent urinary tract infection at the University of Chicago Urogynecology clinic under IRB22-1580.

*E. coli* strains were grown aerobically at 37C in LB broth (BD Difco) while shaking at 250 RPM. Bacteroidaceae strains were routinely grown anaerobically (Coy Lab Products) at 37C with an anaerobic atmosphere of 85% N_2_, 5% H_2_, and 10% CO_2_ in BHIS media: 37g Bacto BHI Premix, 5g Yeast extract (Gibco), 4ml Resazurin (Acros Organics) at 250μg/mL in 1L of DI water before autoclaving, and 10 mL L-cysteine-HCl (EMD Millipore) at 50 μg/mL, 1 mL menadione (MP Biomedical) at 1 mg/mL, and 10 mL hemin (Sigma Aldrich) at 1 mg/mL were added after autoclaving. For colicin kill curve assays, pH modified basal medium (PMBM) was used: 20g Bacto proteose peptone (Gibco), 5g yeast extract (Gibco), and 5g NaCl per 950mL water with pH adjusted to 8.5 with NaOH before autoclaving. Immediately before use, the following was added per 95mL media: 2mL 25% glucose, 2mL 25% K_2_HPO_4_, 1mL 5% L-cysteine, 100μL hemin (5mg/mL), 50μL menadione (5μg/mL), and 5mL 1M sodium bicarbonate. Plates were made with the addition of 15g/L agar (Fisher Scientific).

Antibiotics and other supplements were provided at the following concentrations as noted: 100 μg/mL carbenicillin (Goldbio), 25 μg/mL erythromycin (Sigma), 200 μg/mL gentamicin (Acros Organics). Dilutions of bacteria for CFU assays were performed in 1x phosphate buffered saline (Thermo Fisher Scientific).

### Plasmid Construction and Transformation

Genetic parts used in this chapter are listed in Supplemental Table 2. All parts were cloned into [59]. Genetic parts were amplified using Q5 Hot Start High Fidelity polymerase (New England Biolabs). Genetic parts were then assembled via Gibson assembly. Plasmids were electroporated into E. coli S17 λpir and grown on LB plates with carbenicillin 100 μg/ml. Resulting plasmids were confirmed with Sanger sequencing at the Sanger (DNA) Sequencing Core at the University of Chicago.

### Bacterial Conjugation

To transfer plasmid to Bacteroidaceae strains, 250μL of overnight E. coli culture pelleted at 6,000g for 5 minutes. Pellet was resuspended with 1mL overnight culture of Bacteroidaceae strains and pelleted again. Resulting pellet was resuspended in 20μL BHIS and entire resuspension spotted onto BHIS plate without antibiotics. Plate was incubated overnight aerobically at 37C. The mating spot was then resuspended in BHIS (1mL for *B. thetaiotaomicron, B. ovatus, and P*. vulgatus; 250μL for *B. fragilis*) and 250μL was spread on BHIS plate containing 25 μg/mL erythromycin (Sigma) and 200μg/mL gentamicin (Acros Organics) and incubated for 48 hours anaerobically at 37C. Restreaked, isolated colonies were checked for integration via PCR.

### Uropathogenic *E. coli* Sequencing, Annotation and Analysis

Uropathogenic *E. coli* strains were grown in 5mL of LB overnight at 37C with shaking. Genomic DNA was extracted using DNeasy PowerLyzer Microbial Kit (Qiagen). Bacterial genomes were sequenced and assembled by Plasmidasaurus using Oxford nanopore long reads. Draft genomes were assessed for completeness and contamination using CheckM2 v1.0.2. Genomes passing quality thresholds were annotated using Prokka v1.15.6 with default settings. A pan-genome was constructed using Panaroo v1.6.0, and a core genome alignment was generated with MAFFT as defined by genes present in at least 98% of isolates. Core SNPs were extracted using snp-sites v2.5.1 and used to infer a maximum-likelihood phylogeny with IQ-TREE v3.1 (GTR+G model, 1000 bootstrap replicates). Sequence type, phylogroup, virulence-associated genes, and antimicrobial resistance determinants were identified using the E. coli modules in Kleborate v3.2.4. Multilocus sequence typing (MLST) of *E. coli* strains were determined using MLST 2.0 (https://cge.food.dtu.dk/services/MLST) using the Achtman scheme (Escherichia coli #1) with a minimum allele depth of 5x. Antibiotic resistance was identified using ResFinder 4.7.2 from DTU National Food Institute (https://genepi.food.dtu.dk/resfinder). Virulence factors were identified using VirulenceFinder 2.0.5 (https://cge.food.dtu.dk/services/VirulenceFinder/) using default settings.

### Colicin Soft Agar Overlay Assay

Colicin strains were grown anaerobically overnight in BHIS at 37C. 2.5μl of culture was then spotted onto BHIS agar plate, with multiple peptide-colicins variants spotted on a single plate (1 plate per *E. coli* strain). Colicin strain plates were incubated 16hrs anaerobically at 37C. Resulting spots were removed from the plate by placing cellulose membrane (printer paper) onto the plate with pressure to improve contact. The membrane was removed using tweezers and step was repeated 2-3 times until bacterial spots were no longer visible. Plates were then placed inverted without a lid on a wire baking rack in a glass tray containing paper towels soaked in ∼100mL chloroform. The tray was then covered with cardboard lid and weighted to prevent escape of chloroform vapor and incubated room temperate for 15 minutes. Plates were then removed and allowed to incubate at room temperate with lid cracked to allow chloroform vapor to dissipate and prevent sticking. Top agar was prepared using 0.6% agar in LB, autoclaved, and melted top agar was cooled to 55C in heated bead bath. 15mL culture tubes were preheated in 55C bead bath for 10 minutes. Top agar was distributed into culture tubes (4mL for 100mm circular plate and 5mL 100mm square plate) and maintained in 55C bead bath until immediately before use. Immediately before pouring plates, 5μl of overnight *E. coli* culture was added to top agar, poured atop chloroform-treated plates and rotated to evenly distribute. Plates were then allowed to cool for 20 minutes and incubated overnight at 37C. Susceptibility of *E. coli* to each colicin was rated based upon clearing size (less than, equal to, or greater than initial spot size) and quality (hazy/resistant colonies/full clearing) (Supplemental Figure 2).

### Colicin Kill Curve Assay

For colicin kill curve assays, colicin strains were grown for 16hrs in PMBM anaerobically at 37C. Cultures were then normalized to an OD_600_ of 1 and centrifuged at 4,000g for 10 minutes. Supernatant was then filtered through 0.22μm PES membrane (Nalgene) to remove remaining debris. *E. coli* strains were grown overnight aerobically at 37C in LB with shaking and then subcultured 1:100 until an OD_600_ of 0.2. Cultures were then diluted 1:500 to obtain a concentration of ∼5x10^5^ CFU/mL. 250μL of diluted *E. coli* culture was then added to 250μL of colicin supernatant and incubated aerobically at 37C with shaking. Resulting cultures were plated for CFU dilutions on LB plates at 0, 2, 4, and 24hrs.

### Animal Husbandry

All animal experiments were carried out in compliance with the University of Chicago Institutional Animal Care and Use Committee (IACUC) (Protocol 72610) guidelines and maintained at the University of Chicago Animal Resource Center (ARC). Specific pathogen-free (SPF) mice were purchased from the Jackson Laboratory and maintained in the University of Chicago SPF facility. Mice were housed in cages with paper bedding (ALPHA-dri + PLUS) and given standard chow (Inotiv 2918 Teklad global 18% protein rodent diets, 6% fat). All mice were housed with a 12 hour light/dark cycle at a standard room temperature of 20-24C. All mice were euthanized by CO_2_ asphyxiation followed by cervical dislocation as a secondary measure.

### In Vivo *E. coli* Colonization Prevention by Colicin Secreting *P. vulgatus*

8 week old female C57BL/6 mice were purchased from Jackson Laboratory. Mice were placed on drinking water with 0.625g/L ciprofloxacin and 1g/L metronidazole via water bottle for seven days. Mice were then placed on drinking water with 0.625g/L ciprofloxacin and gavaged with 1mg metronidazole per 1kg body weight to ensure adequate antibiotic dosing (weight was assumed to be ∼20g per mouse). Mice were then returned to regular drinking water for 72 hours to allow for antibiotic washout. Mice were then gavaged with 200μL of ∼10^9^ CFU/mL wild-type or colicin-producing *P. vulgatus* from a fresh culture. Colonization was confirmed by plating on BHIS plates with 25μg/mL gentamicin and 100μg/mL erythromycin after five days with an average titer of 5x10^8^ CFU/g feces (data not shown). Seven days after bacterial gavage, mice were given 5g/L streptomycin in drinking water for 72 hours, followed by a 24 hour washout period. Mice were then gavaged with 200 ul ∼2.5x10^5^ CFU/mL *E. coli* from a fresh overnight culture. Fecal pellets were collected 2, 4, and 7 days after *E. coli* gavage. Fecal pellets were processed by resuspending in 1mL PBS and homogenizing in a Powerlyzer 24 homogenizer (Qiagen) for 2 minutes at 2,000rpm. Samples were then centrifuged at 300g for 30 second. Sample dilutions were plated for CFUs on MacConkey Agar (RPI) plates and incubated overnight at 30C.

### Fecal 16S rRNA Analysis

DNA was extracted from fecal pellets using the QIAmp PowerFecal Pro DNA kit (Qiagen) and submitted to SeqCenter for 16S rRNA sequencing. Samples were prepared using Zymo Research’s Quick-16S kit with phased primers targeting the V3/V4 regions of the 16S gene. Following clean up and normalization, samples were sequenced on a P1 or P2 600cyc NextSeq2000 Flowcell to generate 2x301bp paired end (PE) reads. Quality control and adapter trimming was performed with bcl-convert1 (v4.2.4). Paired-end reads were quality filtered and trimmed (forward: 280 bp; reverse: 220 bp) based on per-base quality scores (>30). Amplicon sequence data were processed in R v4.5.2 using DADA2[60] v.1.38.0 into amplicon sequence variants. Chimeric sequences were removed, taxonomy was assigned using the SILVA database v138.2, and a phyloseq object was constructed using phyloseq v.1.54.2[61]. To account for compositionality, counts were transformed using a centered log-ratio (CLR) transformation after addition of a pseudocount. Aitchison distances (Euclidean distance on CLR-transformed data) were calculated and visualized by principal coordinates analysis (PCoA). Differences in community composition were assessed by permutational multivariate analysis of variance (PERMANOVA) using vegan v2.7.3[62]. Alpha diversity was quantified using the Shannon index. Relative abundances were calculated and taxa aggregated at the genus and family levels for visualization. Differential abundance was assessed using ALDEx2 v.1.42.0[63] with 128 Monte Carlo instances; features present in ≥20% of samples were retained, and p-values were adjusted using the Benjamini–Hochberg false discovery rate. Figures were made in R using ggplot2 v4.0.3.

## Acknowledgments

The authors are supported by the by the National Institute of General Medical Sciences of the National Institutes of Health (R35GM147478; M.M.), by the National Institutes of Health (IMSD Program 5R25GM109439-07, Molecular and Cellular Biology training program T32 GM007183 to J.F.-S.; Research Training in Digestive Diseases and Nutrition T32DK007074-49 to J.G.), by the Arnold and Mabel Beckman Foundation through the Beckman Young Investigator Program (M.M.).

## Figures and Tables

**Supplemental Figure 1.**
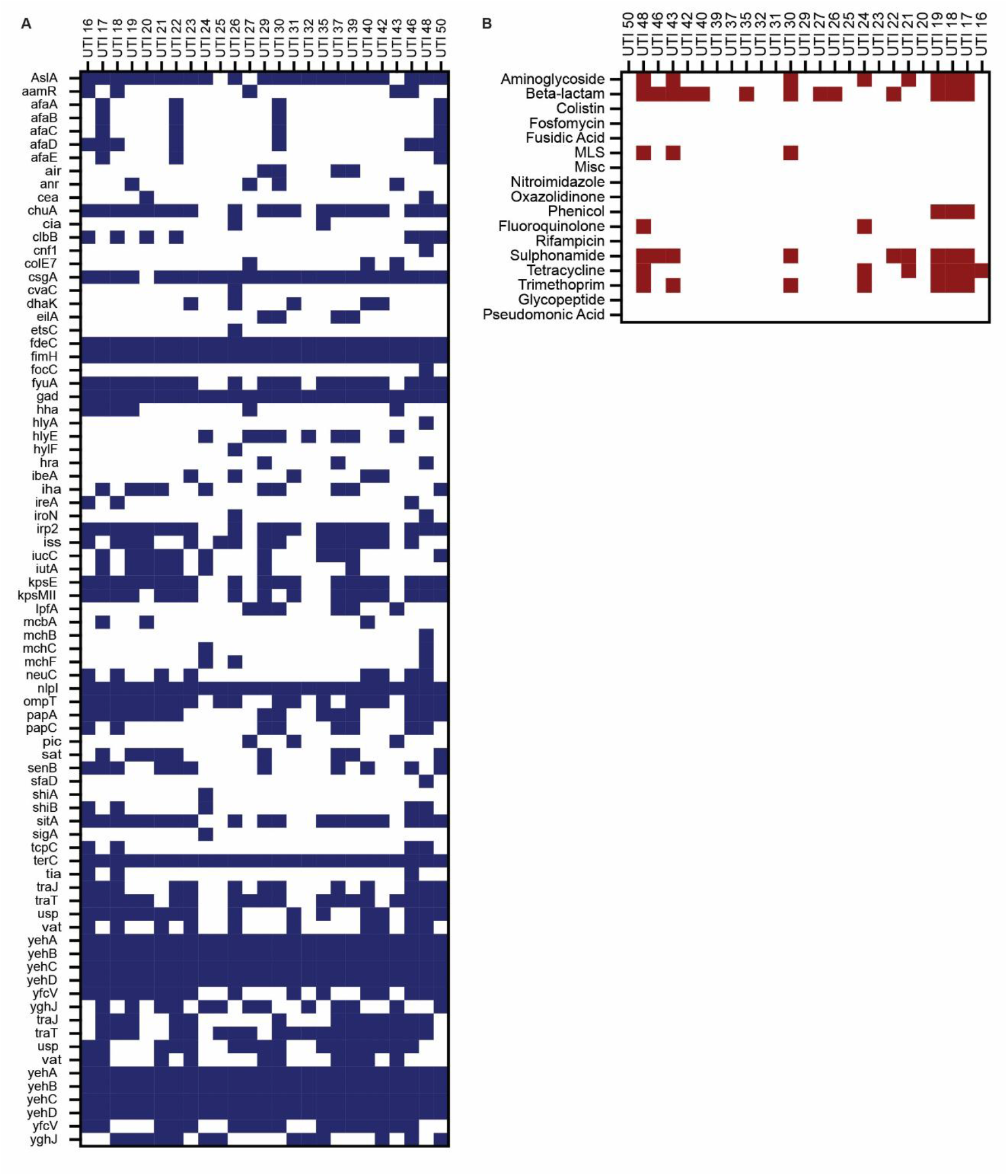
Extended virulence factors and antibiotic resistance markers for primary patient UPEC isolates. (A) Virulence factors were determined for each strain (VirulenceFinder 2.0.5), with blue squares indicating presence of a given gene. (B) . Antibiotic resistance was also identified computationally (ResFinder 4.7.2), with maroon squares indicating resistance for a given antibiotic. (MLS = Macrolides, lincosamides, streptogramins).

**Supplemental Figure 2.**
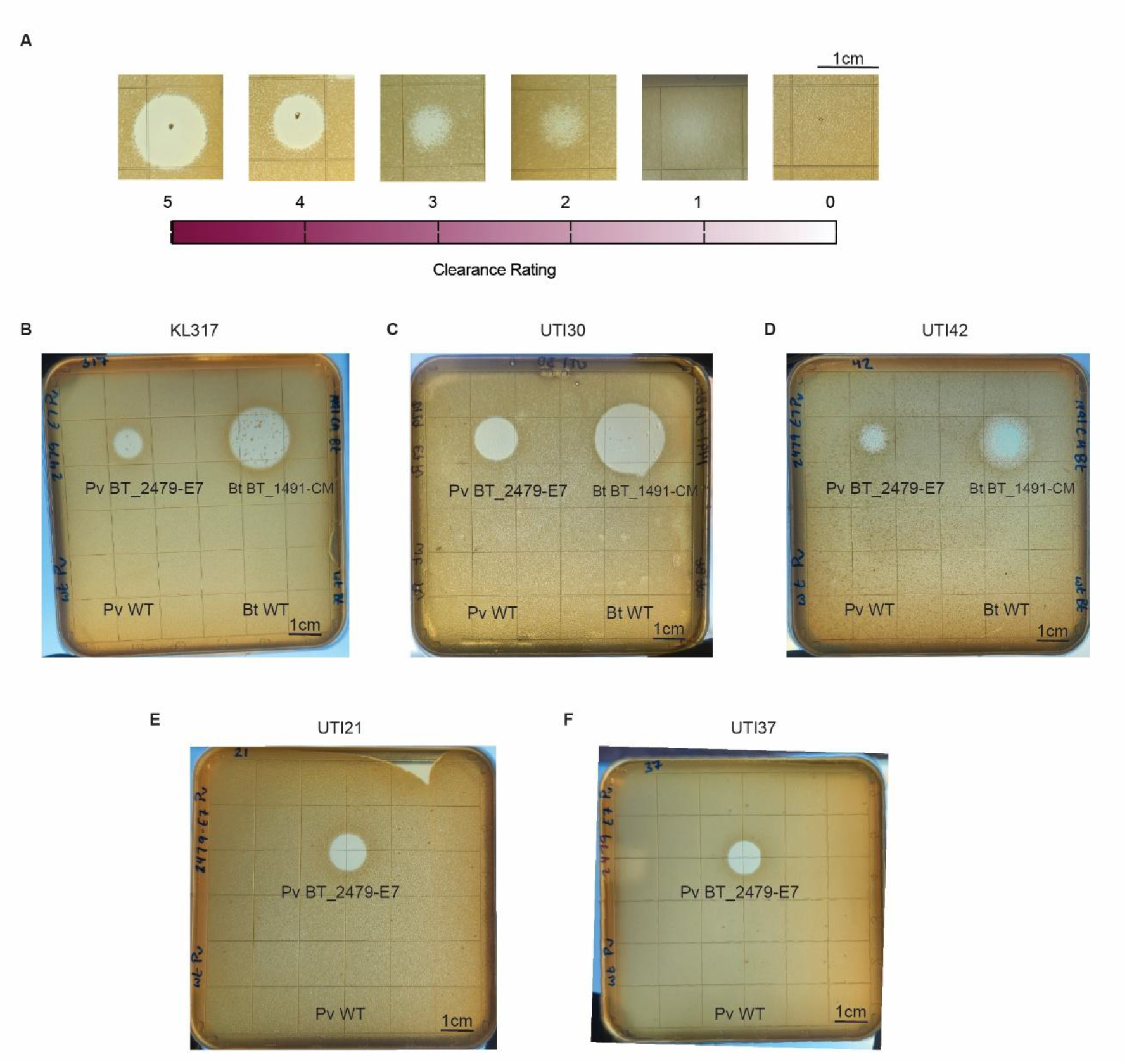
Soft agar overlay assays for determining UPEC susceptibility to colicins secreted by Bacteroidaceae. (A) Representative images of clearance ratings for each spot as follows, with darker shading corresponding to increased clearance: no visible clearance (0), lightened area (1), hazy area (2), clearance smaller than original Bacteroidaceae spot (3), clearance equal to original spot (4), and clearance larger than original spot (5). (B-F) Representative plates for clearance of *E. coli* strains utilized in kill curve assays. For (B) KL317, (C) UTI30, and (D) UTI42, spots were placed as follows: Pv BT_2479-E7 (top left), Bt BT_1491-CM (top right), wildtype *P. vulgatus* (bottom left), and wildtype *B. thetaiotaomicron* (bottom right). For (E) UTI 21 and (F) UTI37, spots were placed as follows: Pv BT_2479-E7 (top) and wildtype *P. vulgatus* (bottom).

